# Arid2 promotes Follicular B-cell differentiation and antibody responses *in vivo*

**DOI:** 10.1101/2025.10.01.678812

**Authors:** Rachel L. Paolini, Sarah Zimmerman, Sadra Marjai, Sofia R. Smith, Pei-Yu Chen, George P. Souroullas

**Author notes:** Address correspondence to GPS George P. Souroullas, PhD, Washington University School of Medicine, 660 S. Euclid Av, St. Louis, MO 63110.

## Abstract

SWI/SNF chromatin remodeling complexes regulate gene expression during development and differentiation. These complexes exist in distinct forms, including the polybromo-associated BAF (PBAF) complex, defined by the ARID2 subunit. ARID2 is frequently mutated in B cell leukemias, but its role in normal B cell development remains unknown. Using conditional knockout mice, we show that Arid2 deletion impairs B cell differentiation *in vivo*. Mb1-Cre–mediated deletion, active in early progenitors, caused a marked reduction of splenic and circulating Follicular B cells, whereas CD19-Cre deletion produced a milder phenotype. These defects were not due to altered proliferation or survival but reflected impaired differentiation potential. Transcriptomic profiling of isolated pro-B, pre-B, and Follicular B cells revealed that Arid2 loss disrupts stage-specific gene expression programs, with cumulative downregulation of B cell receptor signaling and altered lineage-specifying pathways. Functionally, Arid2-deficient mice exhibited impaired germinal center expansion after immunization and reduced IgG antibody production following transplantation. Together, these findings identify Arid2 as a critical regulator of B cell differentiation and maturation by coordinating transcriptional programs during early lymphopoiesis.

## Introduction

B cell development is a multistep process that ensures the generation of a diverse repertoire of mature B lymphocytes that secrete antibodies to fight infection. In the bone marrow, common lymphoid progenitors undergo a series of sequential transitions characterized by the pre-pro-B, pro-B, pre-B, and immature B cell stages to assemble fully functional B cell receptors (BCR) [1]. Non-autoreactive immature B cells differentiate into transitional B cells that migrate to secondary lymphoid organs and mature into Marginal zone B (MZB) or Follicular B cells [1,2]. Upon antigen exposure, MZB and Follicular B cells can become activated via their respective BCRs and mediate the T-cell independent and T-cell dependent immune responses, respectively [1,3]. Each developmental stage is governed by precise signaling and transcriptional networks, and their disruption underlies immunodeficiency, autoimmunity and lymphoid malignancies [4,5].

Epigenetic regulation is central to these processes, controlling lineage-specific gene expression programs and immunoglobulin rearrangements [6]. Key epigenetic regulators include the Polycomb repressive complex 2 (PRC2) [7–9], and SWI/SNF chromatin remodeling complexes [10,11], which alter DNA accessibility to direct gene expression [12,13]. Mammalian SWI/SNF complexes assemble into two main ARID-defined variants: canonical BAF, which contains either ARID1A or ARID1B as mutually exclusive subunits, and polybromo BAF (PBAF), defined by ARID2 [14]. ARIDs are indispensable for complex assembly and their deletion completely inhibits chromatin remodeling activity [14,15]. In B cell neoplasms, ARID1A/1B mutations are common in diffuse-large B cell lymphomas (DLBCL), whereas homozygous deletions in ARID2 occur predominantly in pediatric B-cell acute lymphoblastic leukemias (B-ALL), often originating in early B cell progenitors [16–19]. These patterns suggest distinct roles of BAF and PBAF during B cell lymphopoiesis.

While recent studies have shed light on the function of ARID1A and ARID1B in B cell development [20–22], little is known about ARID2. Here, we used conditional knockout mouse models driven by the B cell-specific Mb1-Cre and CD19-Cre to dissect the stage-specific functions of *Arid2*. We show that Arid2 loss leads to a significant reduction in Follicular B cells, reflecting impaired differentiation rather than altered proliferation or survival. Transcriptomic profiling revealed progressive disruption of BCR-driven programs across development and functional studies demonstrated impaired germinal-center expansion and antibody responses. Together, our findings establish Arid2 as a critical regulator of B cell differentiation and humoral immunity.

## Materials and Methods

### Generation of *Arid2* conditional knockout mice

The *Arid2* mouse strain used for this research project, STOCK *Arid2*^tm1a(EUCOMM)Wtsi^/OuluMmnc, RRID:MMRRC_036982-UNC, was obtained from the Mutant Mouse Resource and Research Center (MMRRC) at University of North Carolina at Chapel Hill, an NIH-funded strain repository, and was donated to the MMRRC by Terry Magnuson, Ph.D., University of North Carolina at Chapel Hill. Mice were generated via IVF and first crossed to *Flpe* mice to generate Arid2^F/+^ mice. Arid2^F/+^ mice were further crossed to heterozygous littermates to generate Arid2^F/F^ mice and then to heterozygous *Mb1-Cre* or *CD19-Cre* mice to generate *Mb1-Cre^+^*, *Arid2^F/F^ Mb1-Cre ^+^*, *CD19-Cre+*, or *Arid2^F/F^ CD19-Cre^+^* mice. All mice were backcrossed to the C57Bl/6 background. Mice used for experiments were aged 4-6 weeks old unless otherwise stated. Allele genotyping was performed on genomic DNA extracted from mouse toe tissue (See **Table 2** for primer sequences). Mice were housed in an Association for Assessment and Accreditation of Laboratory Animal Care (AAALAC)-accredited facility and treated in accordance with protocols approved by the Institutional Animal Care and Use Committee (IACUC) for animal research at Washington University in St. Louis.

### CD19 magnetic enrichment, protein preparation, and immunoblotting

Cells from bone marrow and spleen were pooled from 3-4 mice per group. CD19+ B cells were isolated in accordance with the CD19 microbead protocol (Miltenyi, 130-121-301) using LS columns and MACS separator. CD19+ cells were resuspended in RIPA buffer (Cell signaling) with protease inhibitor. Following sonication, protein was quantified using the DC protein assay (Bio-Rad) and denatured for western blot. 40µg of denatured protein was loaded per sample. Standard western blot procedures were used. Primary antibodies to the following proteins were used: Arid2 (Mouse, Thermofisher, MA5-27862, 1:500) and B-actin (Mouse, Thermofisher, MA5-15739, 1:2000). The secondary antibody was IRDye 680RD Goat anti-Mouse (Licor, 926-68070). Blots were visualized using a LICOR Odyssey Infrared Imaging System.

### Peripheral blood analysis

Peripheral blood was collected into 0.5M potassium EDTA tubes and complete blood counts were counted on a Hemavet (Drew Scientifiic). 2% Dextran in PBS/EDTA was added to each sample, and the top layer containing WBCs were prepped for surface marker staining and flow cytometry. Bone marrow cells from femora and tibiae and spleens/splenocytes were harvested in HBSS buffer supplemented with 2% FBS and 2mM EDTA (HBSS++) using a 26-gauge needle and syringe. Bone marrow and spleen cells were then passed through an 18-gauge needle and a 0.45uM filter. Cells were spun down at ≥500g at 4°C and treated with 1ml of ACK RBC lysis buffer for 5min at room temperature. Cells were washed and resuspended in HBSS++ buffer and kept at 4°C until ready for next steps. For antibody staining, 5-10×10^6^ bone marrow or spleen cells were used per panel. For flow cytometry, unstained, single antibody, and live/dead only controls were used for all experiments. Dead cells were excluded from analysis. The Attune NxT Flow cytometer and FlowJo_v10.10.0 software were used for all flow cytometric analysis. See Tables 1 and 3 for extended antibody information.

### Surface marker antibody staining

For surface marker staining, cells were stained for 30 minutes at 4°C with the appropriate antibodies in HBSS++ buffer. Cells were washed with HBSS++ and resuspended in HBSS++ buffer with propidium iodide and analyzed by flow cytometry (Attune, ThermoFisher). Surface marker antibody cocktails are listed in **Table 3**.

### Annexin-V analysis

For Annexin-V analysis, 10×10^6^ cells were stained for 30 minutes at 4°C with the appropriate surface marker antibodies in HBSS++ buffer. Cells were spun down and resuspended in Annexin-V binding buffer (BioLegend, 422201) and incubated with the Annexin-V antibody (Annexin V-PE/Cyanine7) and 7-AAD for 15 minutes at room temperature before being analyzed by flow cytometry. Surface marker antibody cocktails are listed in **Table 3**.

### Ki67 analysis

For Ki67 analysis, 10×10^6^ cells were stained for 30 minutes at 4°C with the appropriate surface marker antibodies in HBSS++ buffer. Cells were then fixed with Fixation/Permeabilization solution (Cytofix, BD Biosciences) for 20 minutes at 4°C. After fixation, 1×10^6^ of the cells were washed and resuspended in permeabilization buffer (Cytoperm, BD Biosciences) and stained with Ki67 for 30min at room temperature. Cells were washed and resuspended in HBSS++ buffer for analysis.

### Noncompetitive whole bone marrow transplantation

Whole bone marrow was collected from femora and tibiae of Mb1-Cre or Arid2^F/F^ Mb1-Cre donor mice (C57Bl/6, CD45.2) as described above, excluding the ACK RBC lysis step. Whole bone marrow from 3 donor mice were pooled per genotype and then 72×10^6^ cells were washed with HBSS++ buffer and resuspended in 2.4mL of HBSS++ (n+2). Cells were kept on ice until ready for injection. Recipient mice were lethally irradiated with a split dose of 11 Gy, 4-5 hours apart, and 200µL/6×10^6^ of the donor cells were retro-orbitally injected into each recipient mouse the same day (B6, CD45.1). 10 recipient mice were injected per genotype.

### RNA-sequencing on B cell subpopulations

Pro-B, Pre-B, and Follicular B cells were isolated from unimmunized Mb1-Cre and *Arid2*^F/F^ Mb1-Cre mice using fluorescence-activated cell sorting (FACS). The RNAqueous-Micro Total RNA Isolation Kit (Invitrogen) and Purelink RNA Mini Kit (Invitrogen) were used to extract RNA from pro-B cells (≤50K) and pre-B cells (≥500K), respectively. The RNAEasy Plus micro kit (Qiagen) was used to extract RNA from Follicular B cells (≤500K). All samples were DNase treated. Libraries were prepared and sequenced on an Illumina NovaSeq 6000 platform. Differential gene expression analysis was performed using EdgeR and Limma, and pathway analysis utilized GAGE and WGCNA. Washington University Genome Technology Access Center performed library preparation, sequencing, and differential expression and pathway analyses. RNA-seq data are available in Gene Expression Omnibus (GEO), GSE307667.

### Serum Collection and ELISA

Peripheral blood was collected from mice on Day 0 before initial immunization with SRBCs, on Day 21 at re-immunization, and on Day 30. Blood was allowed to coagulate for 1 hour at room temperature, after which it was centrifuged at 10,000 rcf for 10 minutes. The serum was transferred to a clean tube, centrifugation was repeated to ensure complete removal of blood, and the serum was aliquoted and frozen at −80°C. ELISA kits for IgM and total IgG (Invitrogen, 88-50470-86 and 88-50400-22) were used according to the manufacturer’s instructions. Serum was diluted 1:20,000 to 1:80,000 for IgM, and from 1:25,000 to 1:50,000 for IgG. The plates were developed with substrate for 15 minutes before adding stop solution, and then the absorbance was read on a BioTek Synergy HT plate reader at 450 nm.

### SRBC Immunization

For germinal center studies, age-matched mice were immunized intraperitoneally between 8-12 weeks of age with 0.5 ml of a 2% sheep red blood cell (SRBC) suspension in PBS (Cocalico biologicals, 20-1334A). After 21 days, mice were re-injected using the same concentration and volume of SRBC in PBS and sacrificed 9 days later.

### Statistical Analysis

Statistical analysis was performed using GraphPad Prism, excluding RNA-sequencing data. Bar graphs represent mean and standard deviation and student’s t tests were used to compare group differences unless otherwise stated in the figure legend. Sample sizes were determined based on pilot studies or data from similar experiments.

Differential expression analysis of RNA sequencing data was performed with a Benjamini-Hochberg false-discovery rate adjusted p values cut-off of less than or equal to 0.05.

## Results

### Arid2 expression is enriched in early B cell progenitors and Mb1-Cre model achieves most efficient deletion

To better understand the role of Arid2 during B cell development, we first defined its expression in different B cell sub-populations (**Table 4**) using ImmGen datasets [23,24], accessed and visualized via Bloodspot [25]. In mice, *Arid2* is expressed across all B cell subsets, with approximately 4-fold higher expression in pro and pre-B cells than in mature splenic B cells (**Fig. 1a**). This suggested that deleting *Arid2* in early B cell progenitors could reveal both developmental and functional consequences. We therefore used two B cell specific Cre drivers, the Mb1-Cre [26] and CD19-Cre [27] **(Fig. 1b)**. To validate Cre-expression and recombination, we crossed each Cre allele to the Rosa26^LSLtdTomato^ reporter [28]. In the bone marrow, the Mb1-Cre mediated significantly more recombination in pro-B, pre-B, and Immature B cells compared to the CD19-Cre, whereas recombination in Follicular and marginal zone B cells was equally efficient and close to 100% (**Fig. 1c**). In the peripheral blood, TdTomato expression was largely restricted to the B220+ cells, although Mb1-Cre also showed minor activity in CD3+ T cells (**Fig. 1d**). Crossing to the *Arid2* floxed allele (Arid2^F/F^), confirmed complete loss of protein in B cells (**Fig. 1e**). Consistent with gene expression datasets, Arid2 protein expression was significantly higher in bone marrow B cells than the spleen (**Fig. 1a, e**). These data establish that Mb1-Cre more efficiently deletes Arid2 in early progenitors, whereas CD19-Cre-mediated deletion occurs later, a valuable distinction in delineating the function of Arid2 during lymphopoiesis.

**Figure 1.**
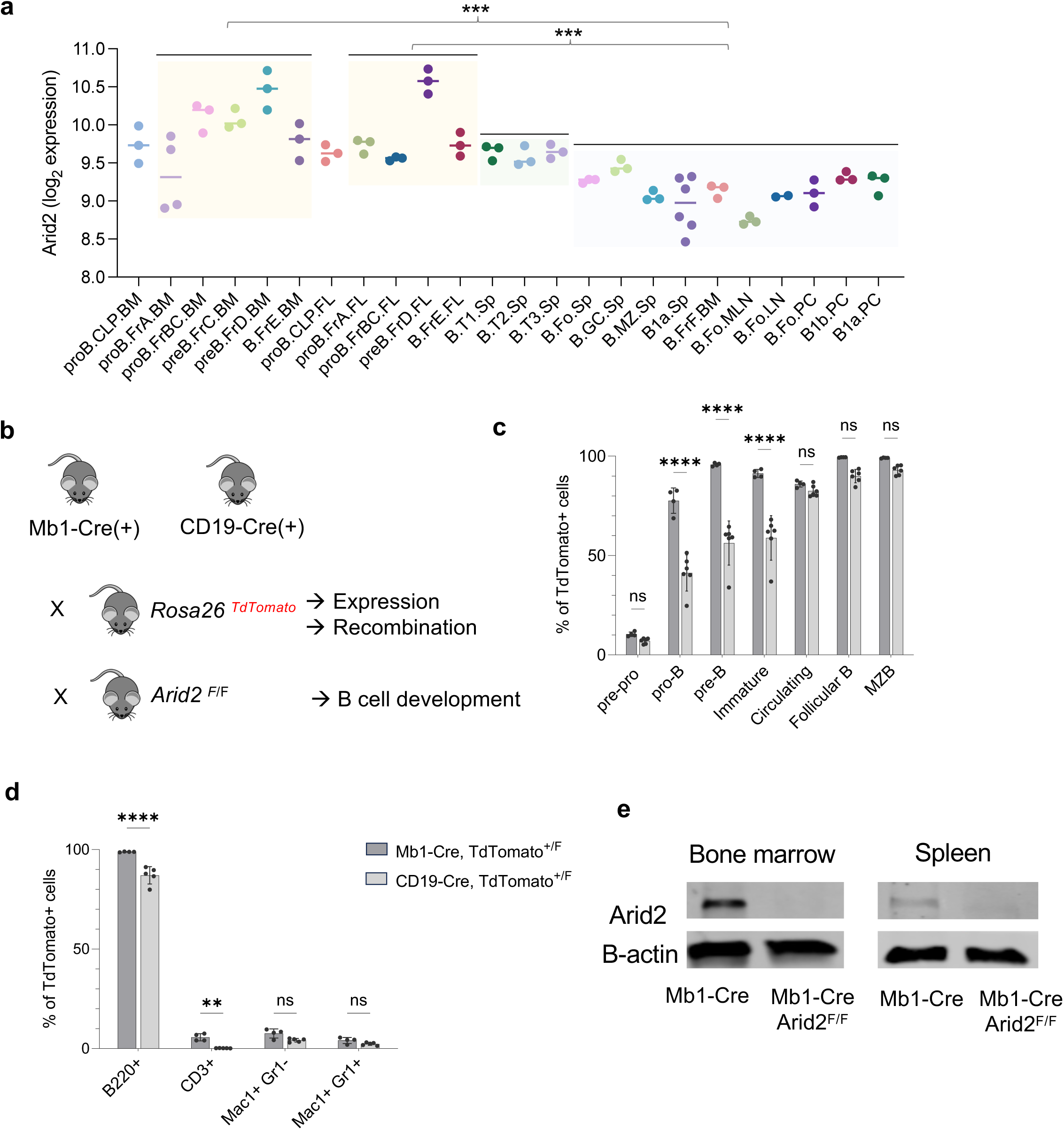
*Arid2* is enriched in early B cell progenitors and efficiently deleted by Mb1-Cre. **a.** Arid2 expression across B cell subsets from ImmGen datasets via Bloodspot. Pro- and pre-B populations in the bone marrow (BM) and fetal liver (FL) where split in two groups, and compared to mature B cell populations from the spleen (SP), lymph node (LN), and the peritoneal cavity (PC). One-way Anova p<0.001. Unpaired t-test of groups as indicated. FoB: Follicular B cells, MZ: Marginal Zone B cells, T1/2: Transitional B cells, CLP: Common Lymphoid Progenitor). *** p<0.0001. Also see **Table 4** for more detailed annotation. **b.** Schematic of TdTomato reporter and conditional knockout strategy using Rosa26^TdTomato^ and *Arid2*^F/F^ alleles crossed to Mb1-Cre or CD19-Cre. **c.** Flow cytometric analysis of TdTomato+ B cells from Mb1-Cre Rosa26^TdTomato/+^ (n=4) and CD19-Cre Rosa26^TdTomato/+^ (n=6). Pre-pro-B, pro-B, pre-B, Immature, and Circulating data from bone marrow, Follicular and MZB data from spleen. **d.** Peripheral blood lineage analysis from the Mb1-Cre and CD19-Cre TdTomato reporter strains. **e.** Western blot showing Arid2 protein expression from CD19+ B cells from bone marrow and spleen of Mb1-Cre mice. Cells were pooled from 3 mice per genotype. *ns* – not significant, **p<0.01, ****p<0.0001 by 2way ANOVA. Error bars indicate standard deviation.

### Early Arid2 deletion via Mb1-Cre results in reduced B cell output and Follicular B cells

We first analyzed peripheral blood by complete blood counts and flow cytometry. Mb1-Cre-mediated *Arid2* deletion resulted in reduced total white blood cells (WBCs) and lymphocytes (**Fig. 2a-b**). Flow cytometry showed that the WBC decrease was driven by fewer B220+ B cells (with a proportional rise in CD3+ T cells) and that absolute B-cell counts were markedly decreased (**Fig. 2c-d**). We did not observe any changes in myeloid populations (**Fig. 2c-d**, **Supp. Fig. 1a-c**).

**Figure 2.**
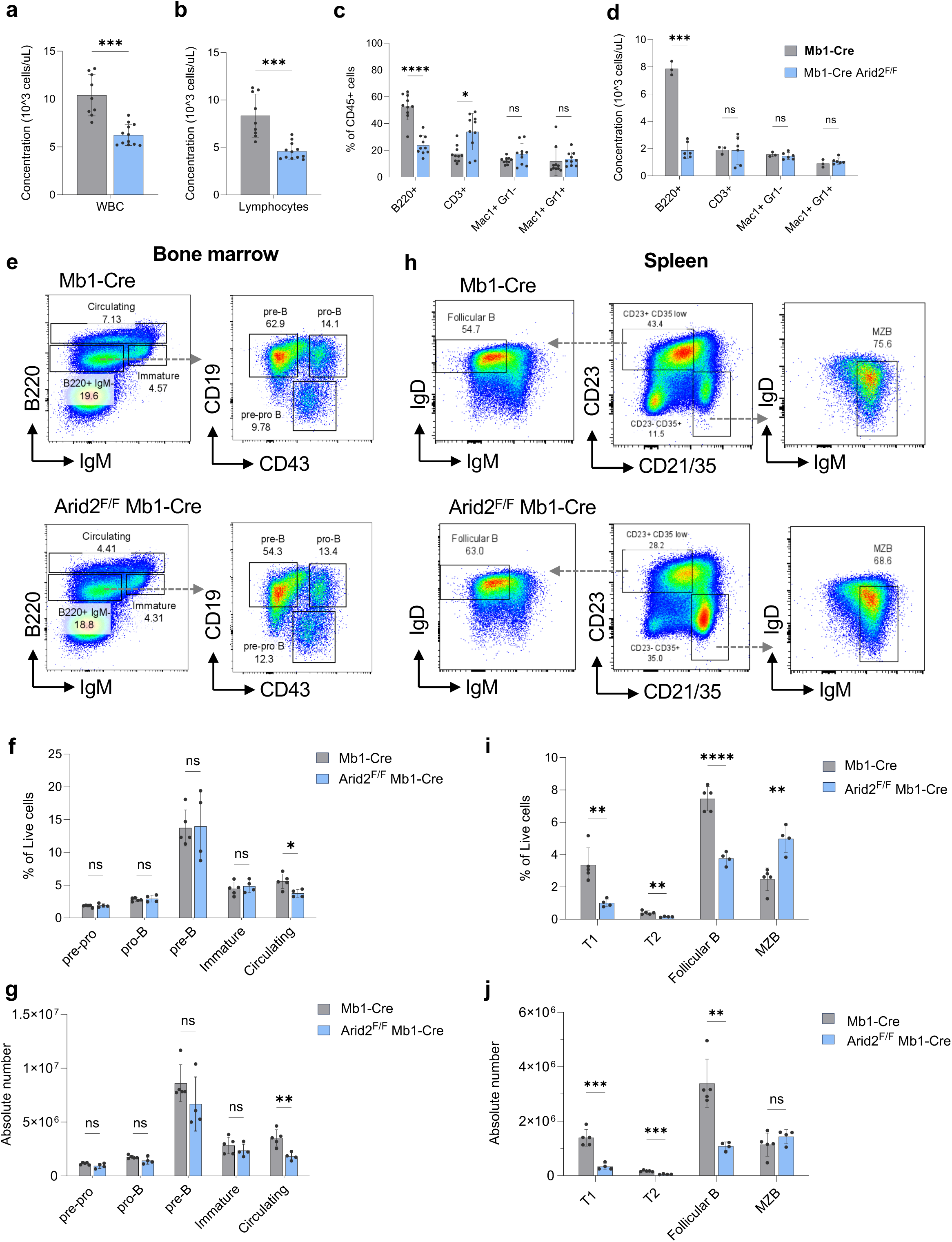
Mb1-Cre–mediated Arid2 deletion reduces peripheral B cell output and Follicular B cells. **a-b.** Complete blood count (CBC) analysis of **a**) white blood cells (WBC) and **b**) lymphocytes in the peripheral blood of Mb1-Cre (n=10) and Arid2^F/F^ Mb1-Cre (n=12) mice. **c.** Flow cytometric analysis of peripheral blood lineage markers (B220+ B cells, CD3+ T cells, Mac1+ Gr1+/- Myeloid cells) as a frequency of CD45+ immune cells from Mb1-Cre (n=10) and Arid2^F/F^ Mb1-Cre (n=10) mice. **d.** Calculated absolute number of B cells, T cells, and myeloid cells in the peripheral blood from white blood cell (WBC) counts (a) and frequency of live cells (c). **e.** Representative flow cytometry plots of B cell populations in the bone marrow. B220 and IgM cells gated from live cells. **f.** Flow cytometric analysis of pre-pro-B, pro-B, pre-B, Immature, and Circulating B cell populations in the bone marrow as a percentage of total live cells. Mb1-Cre (n=5), Arid2^F/F^ Mb1-Cre (n=4). **g.** Absolute number of B cell subsets in the bone marrow described in (f) calculated from total bone marrow counts **h.** Representative flow cytometry plots of Follicular B and MZB cell populations in the spleen. CD23 and CD21/35 cells gated from CD19+, CD43-cells. **i.** Flow cytometric analysis of T1-Transitional (CD19+, CD93+, CD21/35-, CD23-, IgM+, IgD-), T2-Transitional (CD19+, CD93+, CD21/35 (-/low), CD23+, IgM+, IgD+), Follicular B, and MZB cell populations in the spleen as a percentage of total live cells. Mb1-Cre (n=5), Arid2^F/F^ Mb1-Cre (n=4). **j.** Absolute number of populations described in (i) calculated from total spleen counts (Supplementary Figure 2c). *ns* – not significant, *p<0.05, **p<0.01, ***p<0.001, ****p<0.0001 by Welch’s t-test or multiple unpaired t-tests. Error bars indicate standard deviation.

We next examined the bone marrow to determine whether the peripheral B cell phenotype was due to impaired early B cell development. First, we did not find differences in total bone marrow cellularity between Mb1-Cre controls and *Arid2*^F/F^ Mb1-Cre mice (**Supp. Fig. 1g**). Flow cytometry of B cell progenitor populations did not reveal significant changes to the percentage or absolute number of pro-B, pre-B, Immature, or transitional B cells either (**Fig. 2e-g, Supp. Fig. 2a**), indicating intact early development.

In contrast, splenic transitional B cells (T1 and T2), were reduced both in proportion of live cells, but also absolute number in *Arid2^F^*^/F^ Mb1-Cre *Arid2^F^*^/F^ mice compared to controls (**Fig. 2i-j, Supp. Fig. 2c**). Follicular B cells *Arid2*^F/F^ Mb1-Cre were most affected, with an approximately 50% reduction (p <0.0001), corresponding to a loss of 2.3×10^6^ cells (p=0.0038) (**Fig. 2h-j**). Total splenic cellularity was also markedly decreased (**Supp. Fig. 1g**), mainly due to loss of Follicular B cells. Following maturation, naive Follicular B cells migrate between the spleen and bone marrow as part of immune surveillance to respond to infection [29]. In the bone marrow, the circulating B cell population is comprised partially of these cells. Consistent with reduced Follicular B cells in the spleen, *Arid2*^F/F^ Mb1-Cre mice had lower number of circulating B cells in the bone marrow and blood (**Fig. 2e-g**, **Supp. Fig. 2e**). These results identify Follicular B cells as the most sensitive population to loss of Arid2.

### CD19-Cre-mediated *Arid2* deletion results in a milder phenotype

Given the higher expression of *Arid2* in bone marrow progenitors compared to more mature splenic B cells (**Fig. 1a and e**), we hypothesized that earlier deletion with the Mb1-Cre would yield stronger phenotypes than CD19-Cre, which recombines with less efficiency in pro and pre-B cells (**Fig. 1c**). Consistent with this, CD19-Cre-mediated deletion caused more modest changes compared to Mb1-mediated deletion. Total B cell output in the peripheral blood was proportionally decreased, but absolute B cell, WBC and lymphocyte counts were not significantly different from controls (**Fig. 3a-d**). Bone marrow progenitor populations also remained unchanged (**Fig. 3e-g**). In the spleen, while transitional T1 and T2 cells were comparably reduced, the Follicular B cell reduction was substantially smaller in CD19-Cre mice (**Fig. 3h-j**) compared to Mb1-Cre mice (**Fig. 2h-j**). Marginal zone B (MZB) cells increased in frequency (∼60%, p=0.0071) without a change in absolute number (**Fig. 3i-j**) and circulating B cells remained unchanged in the CD19-Cre crosses (**Fig. 3e-g**).

**Figure 3.**
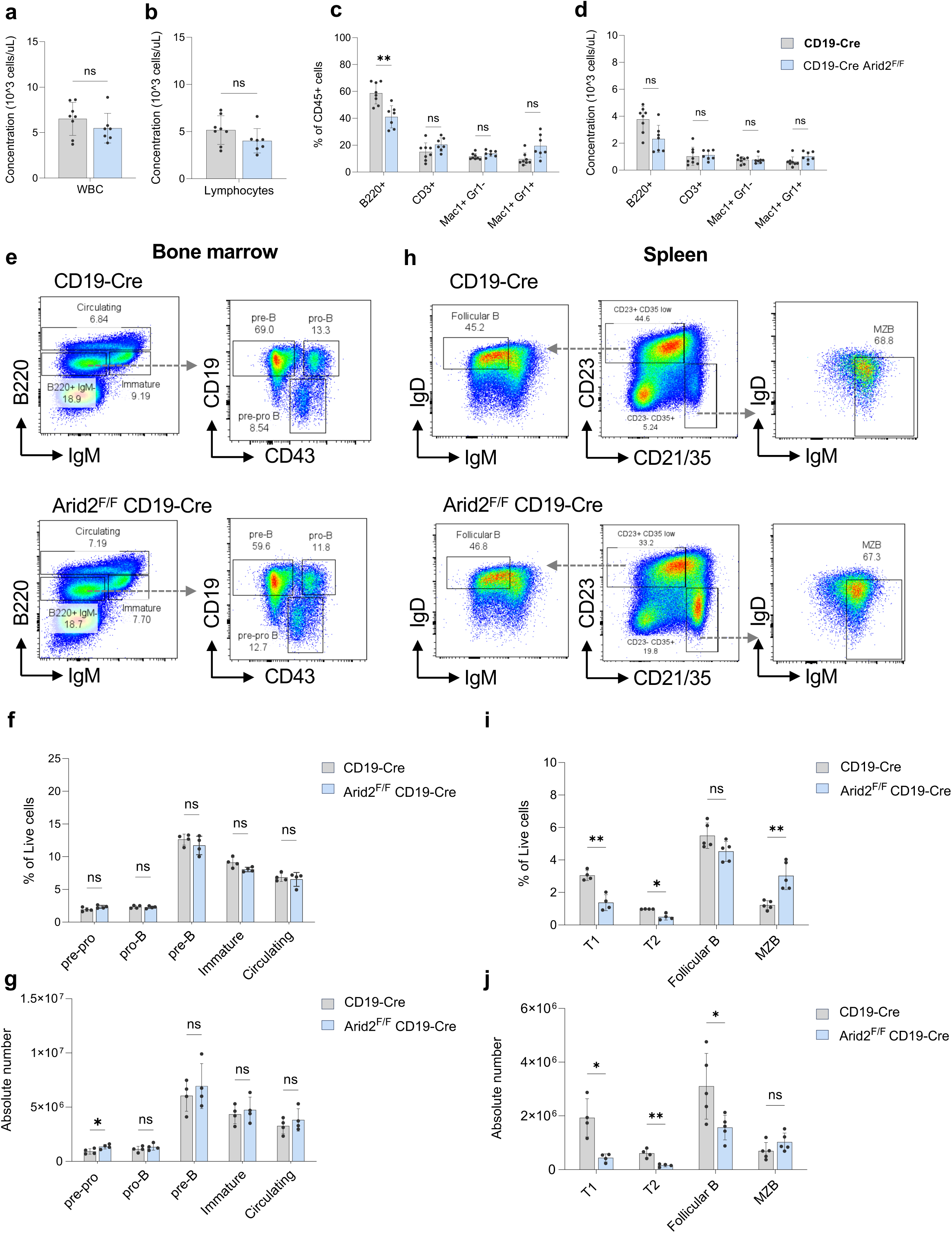
CD19-Cre-mediated Arid2 deletion results in a milder B cell phenotype. **a-b.**Complete blood count (CBC) analysis of **a**) white blood cells (WBC) and **b**) lymphocytes in the peripheral blood of CD19-Cre (n=8) and Arid2^F/F^ CD19-Cre (n=7) mice. **c.** Flow cytometric analysis of peripheral blood lineage markers (B220+ B cells, CD3+ T cells, Mac1+ Gr1+/- Myeloid) as a frequency of CD45+ immune cells from CD19-Cre (n=8) and Arid2^F/F^ Mb1-Cre (n=7) mice. **d.** Calculated absolute number of B cells, T cells, and myeloid cells in the peripheral blood from WBC counts (**c)** and frequency of live cells (**a**). **e.** Representative flow cytometry plots of B cell populations in the bone marrow. B220 and IgM cells gated from live cells. **f.** Flow cytometric analysis of pre-pro-B, pro-B, pre-B, Immature, and Circulating B cell populations in the bone marrow as a percentage of total live cells. CD19-Cre (n=5), *Arid2*^F/F^ CD19-Cre (n=5). **g.** Absolute number of B cell populations described in (**f**) calculated from total bone marrow counts (Supplementary Figure 2d). **h.** Representative flow cytometry plots of Follicular B and MZB cell populations in the spleen. CD23 and CD21/35 cells gated from CD19+, CD43-cells. **i.** Flow cytometric analysis of T1-Transitional (CD19+, CD93+, CD21/35-, CD23-, IgM+, IgD-), T2-Transitional (CD19+, CD93+, CD21/35 (-/low), CD23+, IgM+, IgD+), Follicular B, and MZB cells in the spleen as a percentage of total live cells. CD19-Cre (n=5), Arid2^F/F^ CD19-Cre (n=5). **j.** Absolute number of B cell populations described in (i) calculated from total spleen counts (Supplementary Figure 2d). *ns* – not significant, *p<0.05, **p<0.01 by Welch’s t-test or multiple unpaired t-tests. Error bars indicate standard deviation.

### Arid2 loss impairs B cell differentiation without altering proliferation or survival

The stronger Follicular B cell defect in the Mb1-Cre cohort suggests that the molecular effects caused by the loss of Arid2 are initiated during early B cell development. To test whether reduced Follicular B cells reflect changes in proliferation or survival, we assessed Ki67 and Annexin-V staining, respectively in splenic B cells. We did not find any changes in proliferation nor apoptosis in Arid2-deficient Follicular B or MZB cells (**Fig. 4a and 4b**), suggesting that reduced Follicular B cell numbers were not caused by changes in these processes at steady state.

**Figure 4.**
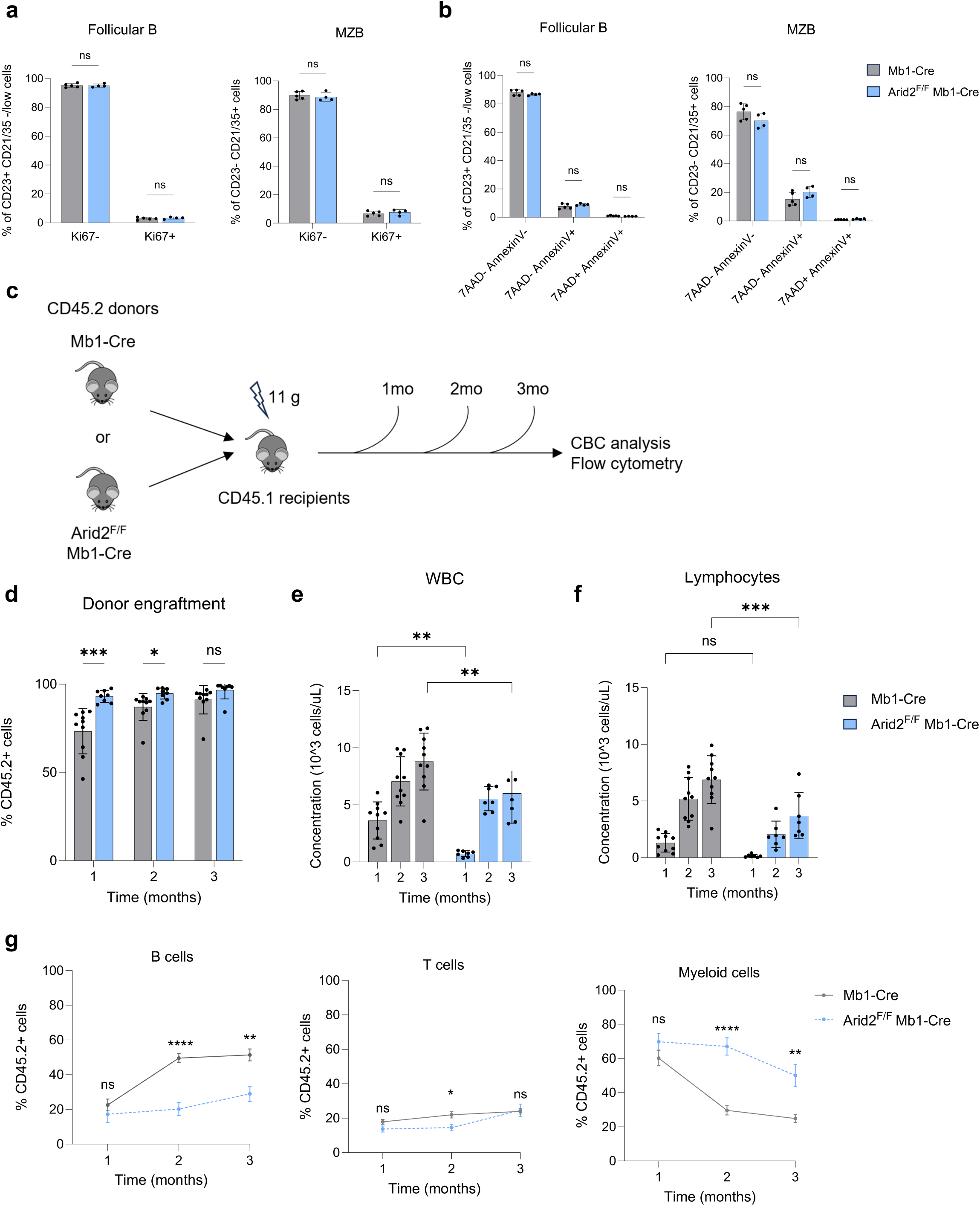
Arid2-deficient progenitors show impaired differentiation but normal proliferation and survival. **a,b.** Flow cytometric analysis of proliferation (Ki67 - panel **a**) and apoptosis (Annexin V – panel **b**) in Follicular B and MZB cells from the spleen of Mb1-Cre (n=5) and *Arid2*^F/F^ Mb1-Cre (n=4) mice. (Viable cells: 7AAD-AnnexinV-, early apoptotic: 7AAD-AnnexinV+, late apoptotic: 7AAD+AnnexinV+). **c.** Schematic of noncompetitive bone marrow transplantation assay. **d.** Flow cytometric analysis of CD45.2+ cells in the peripheral blood of Mb1-Cre (n=10) and Arid2^F/F^ Mb1-Cre (n=7) mice at 1-, 2- and 3-months post-transplant. **e-f.** Concentration of WBC (**e**) and lymphocytes (**f**) in the peripheral blood of recipient mice at 1-, 2- and 3-months post-transplant. Mb1-Cre (n=10) and Arid2^F/F^ Mb1-Cre (n=7). *ns* – not significant, **p<0.01, ***p<0.001, ****p<0.0001 by two-tailed 2-way ANOVA. Error bars indicate standard deviation. **g.** Flow cytometric analysis of peripheral blood lineage markers (B220+ B cells, CD3+ T cells, Mac1+ Myeloid) from Mb1-Cre (n=10) and Arid2^F/F^ Mb1-Cre (n=7) noncompetitive transplant recipient mice as a percentage of CD45.2+ donor cells. *ns* – not significant, *p<0.05, **p<0.01, ***p<0.001, ****p<0.0001 by multiple unpaired t-tests unless otherwise indicated. Error bars indicate standard deviation.

We next tested differentiation potential using bone marrow transplantation, which forces hematopoietic stem cells and progenitors to expand, differentiate, and generate all B cell sub-populations in the host mice. We transplanted whole bone marrow from Mb1-Cre control and *Arid2*^F/F^ Mb1-Cre mice into lethally irradiated recipient mice and analyzed peripheral blood output by flow cytometry and CBC over time (**Fig.4c**). At 1-month, donor engraftment was stably established similarly between groups (**Fig. 4d**). However, total WBC and individual white blood cell subpopulation counts remained generally lower in the recipients of *Arid2-*deficient marrow, specifically due to reduced B cells (**Fig. 4e-g**). Other lineages were largely unaffected. Therefore, in the absence of survival or proliferation defects, the sustained reduction in donor-derived B cells indicates a differentiation impairment intrinsic to Arid2-deficient B cell progenitors.

### Arid2 loss disrupts stage-specific transcriptional programs

To better understand how *Arid2* loss affects B cell differentiation, we performed bulk RNA sequencing on FACS-sorted pro-B, pre-B and Follicular B cells from Mb1-Cre controls and *Arid2*^F/F^ Mb1-Cre mice. Principal component analysis (PCA) showed clear separation between control and knockout samples at all three stages, indicating that transcriptional changes occur as early as the pro-B cell stage (**Fig. 5a**). The number of differentially expressed genes (DEGs) increased progressively (pro-B: 467; pre-B: 715; Follicular B: 2,794), with the large surge in upregulated genes at the Follicular stage (**Fig. 5b**), consistent with cumulative dysregulation. While there was notable overlap in DEGs between adjacent developmental stages such as the pro-B and pre-B cells, as well as between the pre-B and Follicular B cells, a small number of genes was consistently dysregulated across all three groups (**Fig. 5c**), supporting stage-specific regulation.

**Figure 5.**
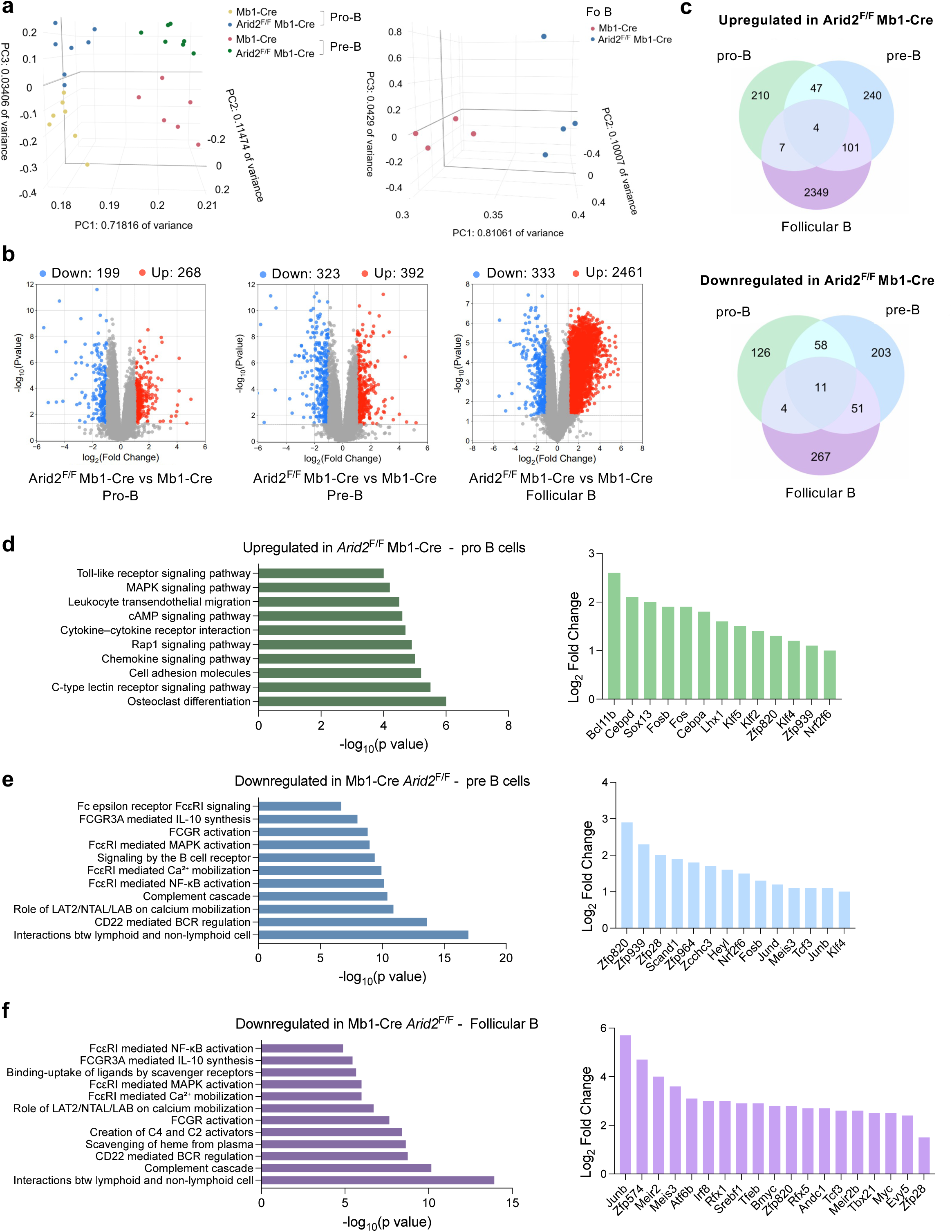
Arid2 loss induces progressive, stage-specific transcriptional dysregulation. **a.** Principal component analysis (PCA) of RNAseq data from sorted pro-B, pre-B and Follicular B cells showing separation between control and knockout samples (n=6-7 mice per group) **b.** Volcano plots of differentially expressed genes (DEG) from pro-B (left), pre-B (middle), and Follicular B (right) data. Significantly upregulated (1≤) and downregulated genes (≤-1) are red and blue, respectively. adj-p≤0.05, and FDR≤0.05. **c.** Venn Diagram depicting overlap in DEG between the Arid2-deficient pro-B, pre-B, and Follicular B cells. Arid2^F/F^ Mb1-Cre pro-B (green), Arid2^F/F^ Mb1-Cre pre-B (blue), and Arid2^F/F^ Mb1-Cre Follicular B (purple), correspond to colors in e. **d-f.** Selected enriched gene sets important for B cell differentiation (left), and fold change of related transcription factors (right) in (**d**) *Arid2*^F/F^ Mb1-Cre pro-B cells (KEGG signaling and metabolism database), (**e**) *Arid2*^F/F^ Mb1-Cre pre-B (Canonical pathways, Reactome database), and (**f**) *Arid2*^F/F^ Mb1-Cre Follicular B cells (Canonical pathways, Reactome database). adj-p≤0.05, and FDR≤0.05. **e.** Fold change of transcription factors or co-factor genes within Arid2-deficient pro-B, pre-B, or Follicular B cells.

Pathway enrichment analysis of Arid2-deficient pro-B cells, using the Generally Applicable Gene-set Enrichment method (GAGE) [30], identified upregulation of pathways related to osteoclast differentiation, cytokine signaling, cell adhesion/migration, and toll-like receptor signaling (**Fig. 5d**); processes that influence early lineage commitment [31–33]. These changes were accompanied by increased expression of transcription factors known to regulate these pathways, including *Cebpa*, *Cebpd, and* AP-1 members *Fos* and *Fosb* (**Fig. 5d**). Pre-B cells showed continued AP-1 activation (Jun, Jund, Junb), linking early and intermediate transcriptional changes (**Fig.5e**). In contrast, B cell receptor (BCR) signaling and Fc epsilon Receptor I (FcεRI)- mediated MAPK, Calcium, and NF-kB pathways were significantly downregulated in Arid2-deficient pre-B and Follicular B cells (**Fig. 5f**). This aligns with the Follicular vs MZB shift observed in Arid2-deficient mice, as stronger BCR signaling favors Follicular fate whereas weaker signaling favors MZB [34,35].

### Arid2-deficient mice retain IgG production at steady state despite smaller germinal centers

Because BCR signaling underlies mature B cell activation, we hypothesized that the downregulation of BCR signaling in Arid2-deficient mice could lead to defective B cell activation and antibody production in *Arid2*^F/F^ Mb1-Cre mice. To test this hypothesis, we analyzed IgM and IgG antibody concentrations in the serum of unimmunized Mb1-Cre and *Arid2*^F/F^ Mb1-Cre mice and mice immunized with sheep red blood cells (SRBCs) (**Fig. 6a**). ELISA of serum from unimmunized mice (day 0) showed a two-fold increase in IgM, consistent with more MZB cells, but no change in IgG despite reduced Follicular B cells (**Fig. 6b-d**). This suggests that Follicular B cells from Arid2-deficient cells retain the ability to be activated upon antigen recognition. Transcriptomic analysis also supports the above results. Specifically, Arid2-deficient Follicular B cells upregulated genes in JAK-STAT, Ras, and ER stress pathways (e.g. *Jak3, Stat3, Socs3, Junb, Irf8, Myc*) (**Supp. Fig. 4f**), programs linked to germinal center activity and plasma cell differentiation [36–38], and may explain how Arid2-deficient Follicular B cells overcome reduced BCR signaling to elicit a T-cell dependent immune response.

**Figure 6.**
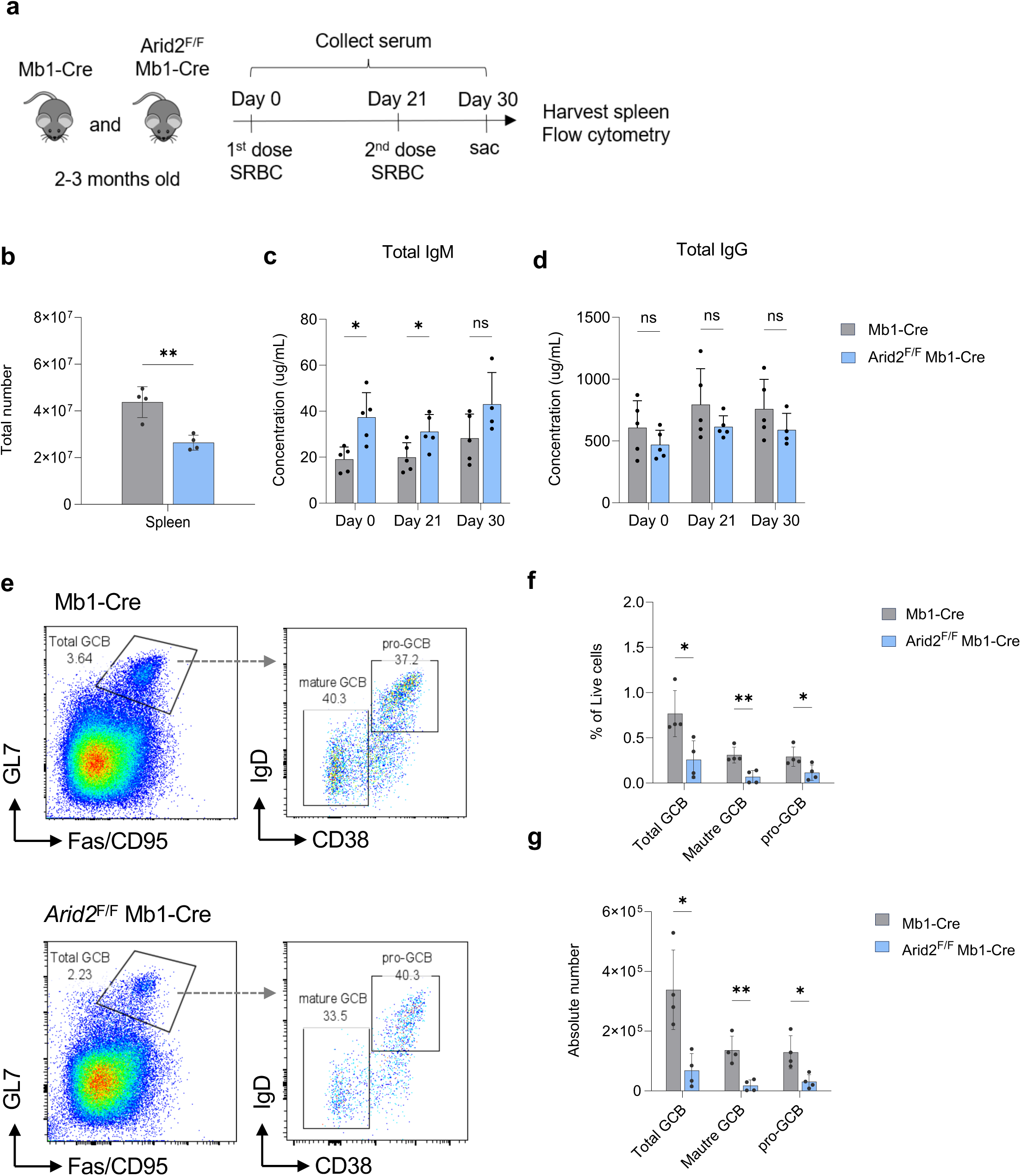
Arid2-deficient mice exhibit altered antibody responses to immunization. **a.**Schematic of the SRBC immunization protocol **b.** Total spleen cell counts in mice of indicated genotypes **c-d.** ELISA measurements of IgM (**b**) and IgG (**c**) antibody concentrations from the peripheral blood of immunized Mb1-Cre and *Arid2*^F/F^ Mb1-Cre mice. **e.** Representative flow cytometry plots of GCB cells from spleens of immunized Mb1-Cre and *Arid2*^F/F^ Mb1-Cre mice. **f.** Flow cytometric analysis of GCB cells in the spleen of immunized mice as a percentage of total live cells. **g.** Absolute number of GCB cells described in (**e**) calculated from total spleen counts (b). *ns* – not significant, *p<0.05, **p<0.01 by Welch’s t-test or multiple unpaired t-tests. Error bars indicate standard deviation.

In immunized mice, as expected, the concentration of serum IgM at Day 21 remained significantly higher in the Arid2-deficient mice (**Fig. 6c**), consistent with persistent MZB activity. Next, we used flow cytometry to analyze the germinal center response in the spleen, a critical component of the T-cell dependent immune response and IgG production. We found significantly fewer GCB cells (pro GCB and mature GCB populations) (**Fig. 6e-g),** in contrast to the fact that IgG levels were maintained (**Fig. 6d**). These data suggest that despite reduced germinal center size, Arid2-deficient Follicular B cells can still seed a germinal center reaction and generate IgG-secreting plasma cells.

### Transplantation stress unmasks intrinsic defects in T-cell dependent IgG responses

To determine whether the observed phenotypes reflect cell intrinsic B cell defects and not niche effects, we transplanted bone marrow from Mb1-Cre controls or *Arid2*^F/F^ Mb1-Cre mice into irradiated recipients and immunized them with SRBC four months later when immune populations had stabilized. Thirty days after immunization, recipient mice of *Arid2*^F/F^ Mb1-Cre bone marrow had significantly lower spleen cellularity and weight compared to mice transplanted with Mb1-Cre- only controls (**Fig. 7a, Supp. Fig. 4**), consistent with reduced B cell output. At Days 0, 21, and 30 after immunization, the concentration of serum IgM was similar between the two groups (**Fig. 7b**). However, serum IgG levels were significantly lower in mice transplanted with *Arid2*^F/F^ Mb1-Cre bone marrow compared to those transplanted with control (**Fig. 7c**). Flow cytometry confirmed significantly fewer pro- and mature germinal center B cells after immunization (**Fig. 7d-f**). Thus, removing potential niche influences and adding transplant stress revealed that Arid2-deficient progenitors have an intrinsic defect in T-cell dependent antibody responses.

**Figure 7.**
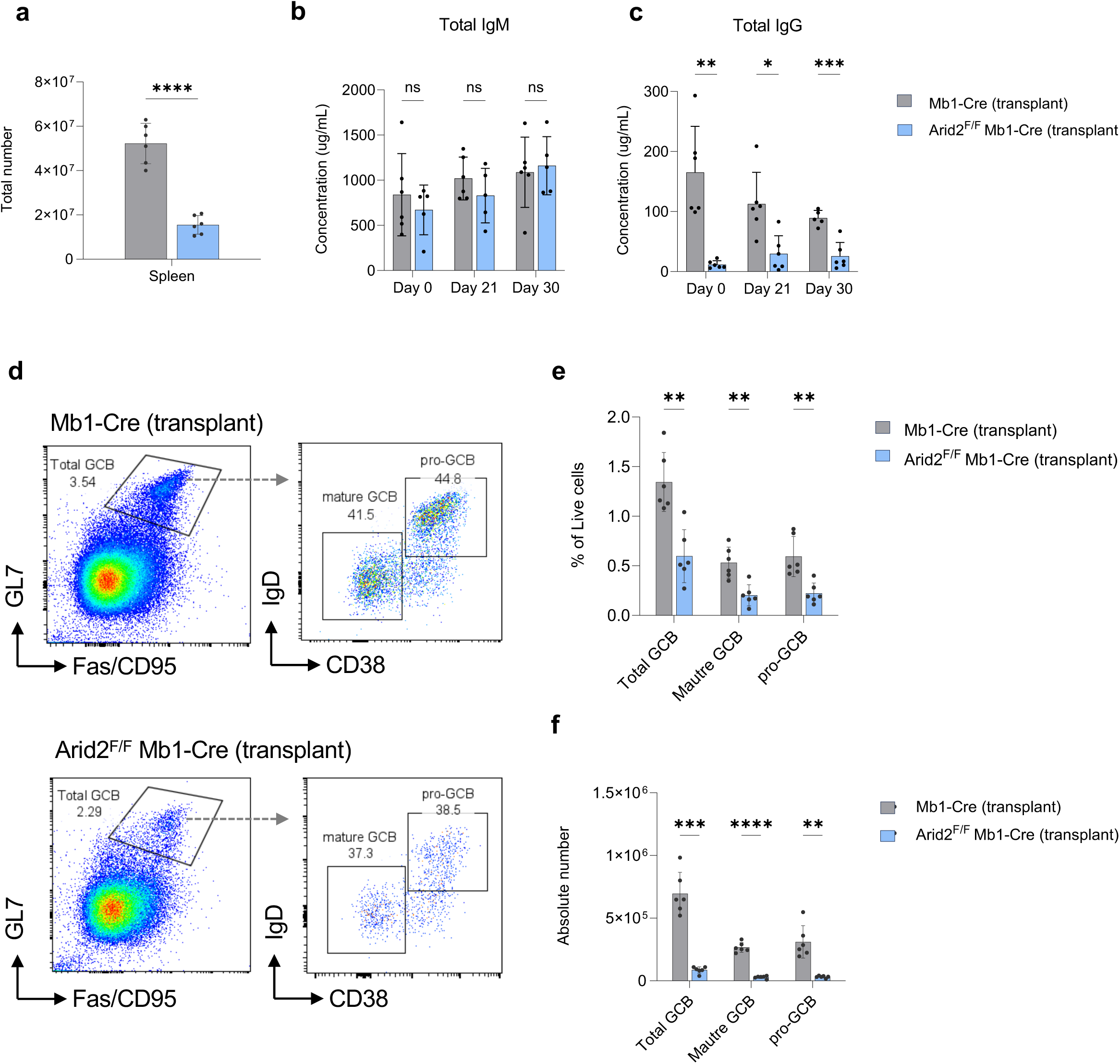
Transplantation stress unmasks intrinsic defects in germinal center and IgG responses. **a.**Reduced spleen weight and cellularity in transplant recipients of *Arid2*^F/F^ Mb1-Cre bone marrow after SRBC immunization. (n=5) cells. **b.** Serum IgM concentrations of recipients of *Arid2*^F/F^ Mb1-Cre vs Mb1-Cre control bone marrow. **c.** Serum IgG concentrations significant reduced in recipients of *Arid2*^F/F^ Mb1-Cre marrow. **d.** Representative flow cytometry plots of GCB cells from spleens of immunized mice transplanted with Mb1-Cre and *Arid2*^F/F^ Mb1-Cre cells. **e.** Flow cytometric analysis of GCB cells in the spleen of immunized transplant mice as a percentage of total live cells. **f.** Absolute number of GCB cells described in (e) calculated from total spleen counts (a). *ns* – not significant, *p<0.05, **p<0.01, ***p<0.001, ****p<0.0001 by Welch’s t-test or multiple unpaired t-tests. Error bars indicate standard deviation.

## Discussion

In this study, we identify Arid2 as a stage-specific regulator of B cell development. Using two complementary Cre models, we demonstrate that the timing of *Arid2* deletion determines the severity of the B cell phenotype. More efficient deletion in early B cell development by the Mb1-Cre resulted in a more significant reduction in splenic and circulating Follicular B cells compared to the CD19-Cre, which causes a milder phenotype. Normal numbers of bone marrow B cell progenitors suggest that Arid2 is not required for early B cell survival or expansion but instead is critical for differentiation into mature B cells.

Transcriptomic analysis by RNAseq revealed that Arid2 loss alters transcriptional programs progressively across development. Upregulation of AP-1 transcription factors (*Fos, Fosb, Jun* family) and cytokine/adhesion pathways appeared in pro-B cell progenitors, while BCR and FCεRI signaling were suppressed in pre-B and Follicular B cells. This pattern of expression suggests that Arid2 normally restrains early transcriptional networks that interfere with B cell maturation and instead promotes signaling required for Follicular B cell fate. These findings are consistent with prior work, where SWI/SNF complexes cooperate with AP1 in other cell types to regulate differentiation [39], and their imbalance may contribute to the observed reduction in Follicular B cells and relative expansion of MZB cells. AP1 members can also regulate CD40-mediated B cell activation [40] and plasma cell differentiation [41], which could explain the ability of Arid2-deficient Follicular B cells to remain activated despite lower number of germinal center B cells.

Functionally, Arid2-deficient Follicular B cells still seeded germinal centers and produced IgG at steady state, despite reduced GC size. This indicates the presence of compensatory transcriptional programs, such as JAK-STAT and ER stress responses, which can support plasma cell differentiation and antibody production. In contrast, bone marrow transplantation revealed a more severe phenotype, with reduced spleen cellularity, fewer GC B cells, and significantly diminished IgG titers. This difference likely reflects two independent variables. First, the stress of reconstitution places greater demand on B cell progenitors, unmasking the differentiation defect more clearly, and second, the transplantation model controls for possible niche effects, revealing cell intrinsic defects of Arid2-deficient B cells.

Our study underscores the importance of Arid2 in safeguarding lineage fidelity and sustaining humoral immunity. It also raises new questions, such as whether antibodies secreted from Arid2-deficient cells are fully functional, whether chronic Ras/AP-1 signaling predisposes cells to autoreactivity as previously suggested in a different setting [42], and how stress pathways in Follicular B cells contribute to potential functional compensation. Given the prevalence of mutations in SWI/SNF components in human cancers [43], our work also raises the possibility that loss of Arid2 both compromises normal immune responses and primes cells for malignant transformation. Future studies should define how Arid2 cooperates with other lineage regulators and explore how chromatin accessibility changes at specific loci within the context of broader epigenetic regulation of B cell differentiation.

## Acknowledgements

We thank the Siteman Flow Cytometry Core for flow cytometry services, the McDonnell Genome Institute/Genome Technology Access Center at Washington University for RNA-sequencing and genome analysis, and the Department of Comparative Medicine for animal maintenance and expertise. We are grateful to members of the Souroullas lab for insightful discussions and critical input on data analysis and presentation. We also thank the Bednarski lab (Washington University in St. Louis) for generously sharing reagents and protocols and for expert guidance on B-cell biology. The Siteman Cancer Center is supported in part by an NCI Cancer Center Support Grant #P30CA091842. Rachel Paolini was partially supported by an NIH NHGRI Institutional Training Grant in Genomic Science (T32HG000045). The content is solely the responsibility of the authors and does not necessarily represent the official views of the NIH.

## Author contributions

RLP, SZ, SM, SRS, PYC and GPS performed experiments and analyzed data. RLP and GPS conceived the experimental design and wrote the manuscript. GPS acquired funding and supervised the project.

## Notes

https://www.ncbi.nlm.nih.gov/geo/query/acc.cgi?acc=GSE307667

### Competing Interest Statement

The authors have declared no competing interest.

