## Supplemental Figure Legends for "Arid2 promotes Follicular B-cell differentiation and antibody responses *in vivo*"

**Supp. Figure 1. Immune cell distribution in *Arid2* conditional knockout mice.**

**a.** Representative flow cytometry plots of B cells, T cells, and myeloid populations in the peripheral blood of *Arid2*$^{F/F}$ Mb1-Cre and Mb1-Cre control mice.
**b-c.** Concentration of blood populations (**b**) and percent of white blood cells (**c**) in the peripheral blood of *Arid2*$^{F/F}$ Mb1-Cre and Mb1-Cre control mice.
**d**. Representative flow cytometry plots of B cells, T cells, and myeloid populations in the peripheral blood of *Arid2*$^{F/F}$ CD19-Cre and CD19-control mice.
**e-f.** Concentration of blood populations (**b**) and percent of white blood cells (**c**) in the peripheral blood of *Arid2*$^{F/F}$ CD19-Cre and CD19-Cre control mice.
**g**. Total number of bone marrow and spleen cells from Mb1-Cre (n=10) and *Arid2*$^{F/F}$ Mb1-Cre (n=7) mice.

**h**. Total number of bone marrow and spleen cells from CD19-Cre (n=9) and *Arid2*$^{F/F}$ CD19-Cre (n=9) mice.

*ns* – not significant, **p<0.01, ***p<0.001, ****p<0.0001 by Welch's t-test or multiple unpaired t-tests. Error bars indicate standard deviation.

**Supp. Figure 2. Follicular and Transitional B cell counts in the bone marrow and peripheral blood of Arid2 conditional knockout mice**

**a.** Percentage (left panel) and Absolute number (right panel) of Transitional B cells in the bone marrow of Mb1-Cre (n=5) and *Arid2*$^{F/F}$ Mb1-Cre (n=4) mice.

**b.** Percentage (left panel) and Absolute number (right panel) of Transitional B cells in the peripheral blood of Mb1-Cre (n=5) and *Arid2*$^{F/F}$ Mb1-Cre (n=5) mice.

**c.** Representative flow cytometry plots of Transitional B cells in the spleen of Mb1-Cre and *Arid2*$^{F/F}$ Mb1-Cre mice (in context to main Fig. 2)

**d.** Percentage (left panel) and absolute number (right panel) of Follicular B cells in the bone marrow of Mb1-Cre (n=5) and *Arid2*$^{F/F}$ Mb1-Cre (n=4) mice.

**e.** Percentage (left panel) and Absolute number (right panel) of Transitional B cells in the peripheral blood of Mb1-Cre (n=6) and *Arid2*$^{F/F}$ Mb1-Cre (n=6) mice.

**f.** Percentage (left panel) and Absolute number (right panel) of Follicular B cells in the bone marrow of CD19-Cre (n=4) and *Arid2*$^{F/F}$ CD19-Cre (n=4) mice.

**g.** Percentage (left panel) and Absolute number (right panel) of Transitional B cells in the peripheral blood of CD19-Cre (n=4) and *Arid2*$^{F/F}$ CD19-Cre (n=4) mice.

*ns* – not significant, *p<0.05, **p<0.01, ***p<0.001, ****p<0.0001 by multiple unpaired t-tests. Error bars indicate standard deviation.

**Supp. Figure 3. Extended RNA-seq analysis: transcription factor dysregulation and pathway enrichment in Arid2-deficient B cells.**

**a-c.** Hierarchical clustering of all differentially expressed genes in Arid2-deficient pro-B (a), pre-B (b) and Follicular B cells (c).

**d.** Upregulated and downregulated genes in *Arid2*$^{F/F}$ Mb1-Cre **pro-B** cells related to B cell differentiation and cell fate.

**e.** Upregulated and downregulated genes in *Arid2*$^{F/F}$ Mb1-Cre **pre-B** cells related to B cell differentiation and interferon signaling.

**f.** Upregulated and downregulated genes in *Arid2*$^{F/F}$ Mb1-Cre **Follicular B cells** related to B cell differentiation and mature B cell activation.

**Supp. Figure 4. Spleen weight and appearance in *Arid2* conditional knockout mice**

**a-b.** Spleen weights (a) and appearance (b) of *Arid2*$^{F/F}$ Mb1-Cre and Mb1-Cre control mice, and mice transplanted with bone marrow from these animals.

*ns* – not significant, *p<0.05, **p<0.01, ***p<0.001, ****p<0.0001 by two-tailed 2-way ANOVA. Error bars indicate standard deviation.
