## Supplemental Figures for "Arid2 promotes Follicular B-cell differentiation and antibody responses *in vivo*"

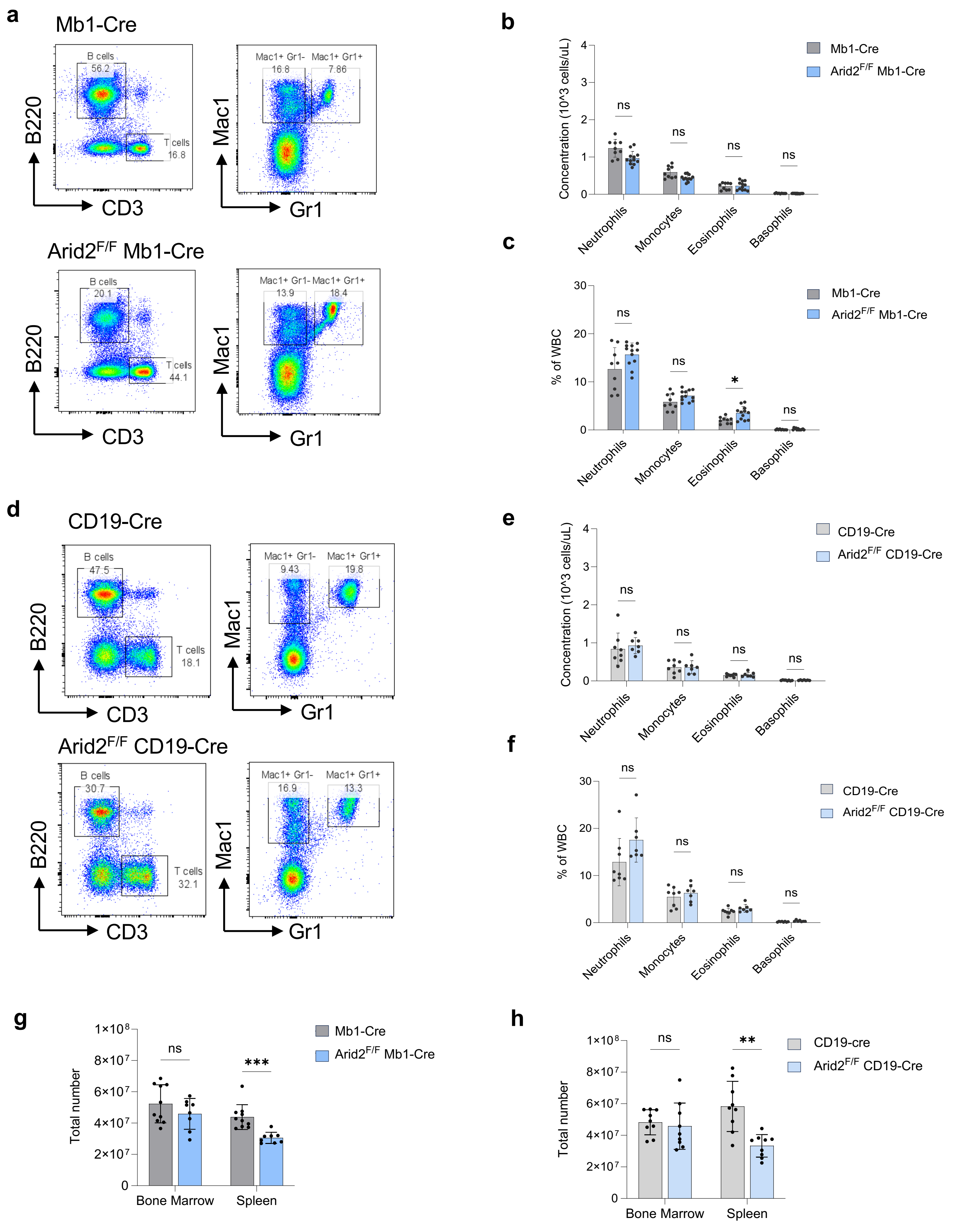


**Figure S1.** Immune cell distribution in *Arid2* conditional knockout mice.

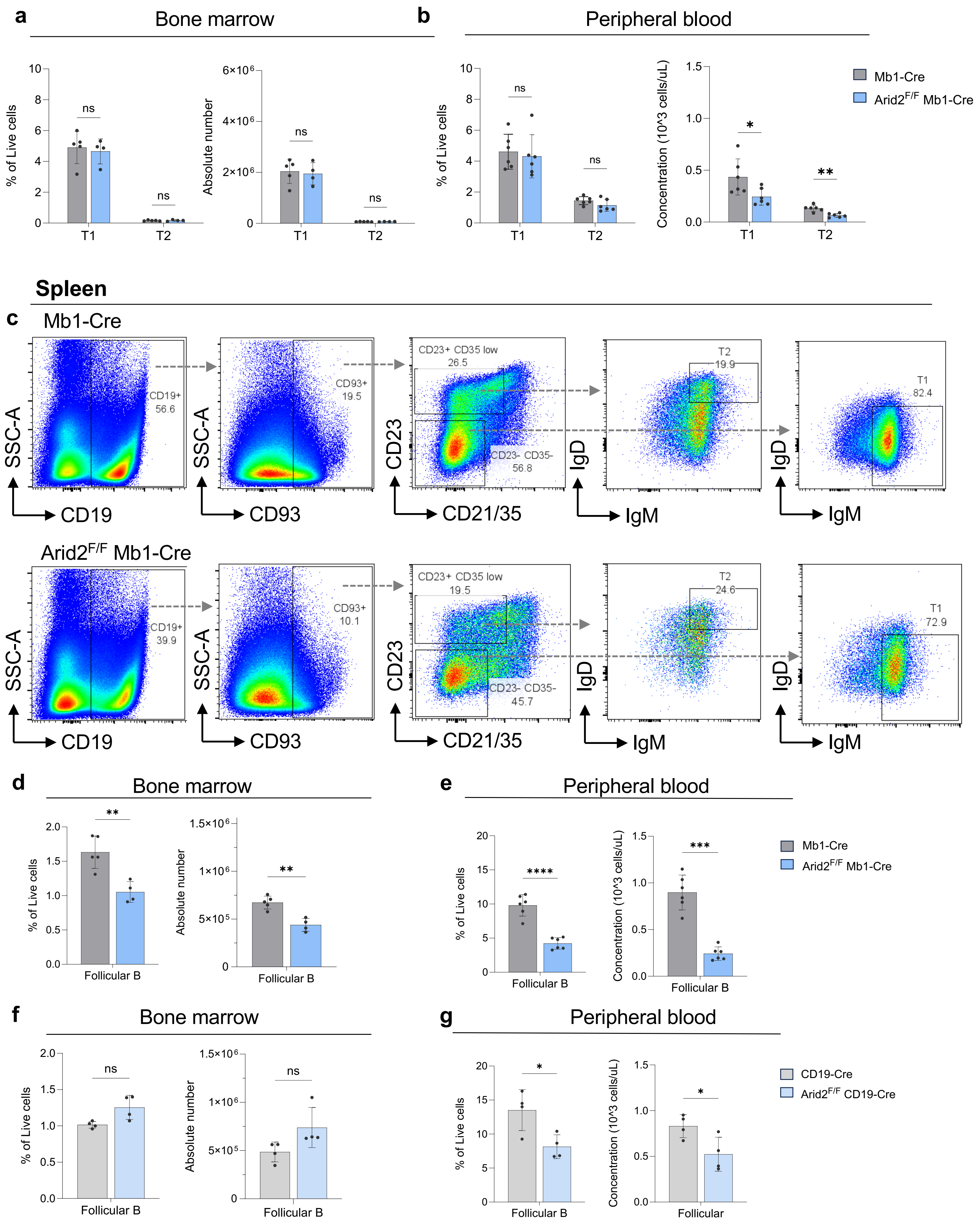


**Figure S2.** Follicular and Transitional B cell counts in the bone marrow and peripheral blood of *Arid2* conditional knockout mice

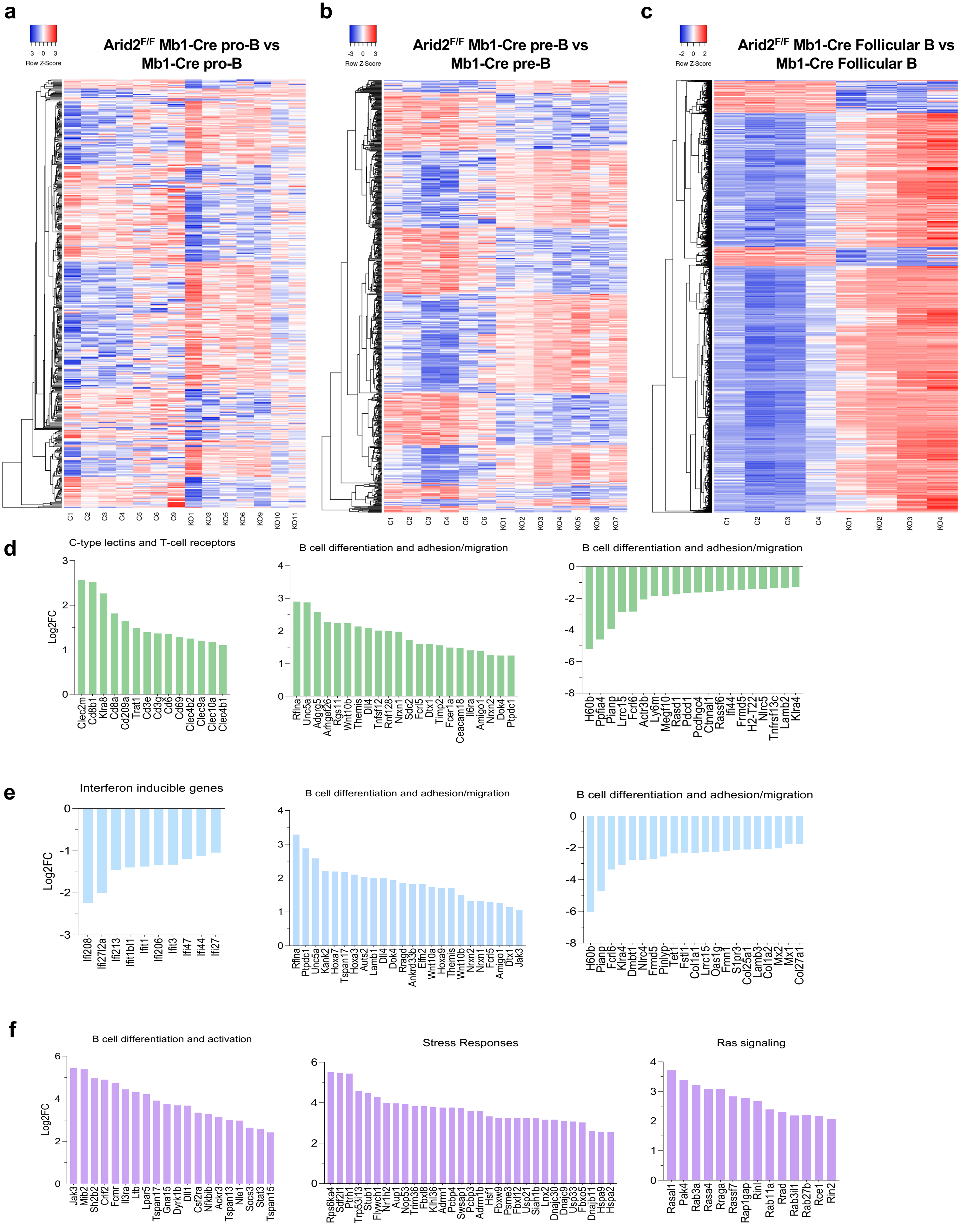


**Figure S3.** Extended RNA-seq analysis: transcription factor dysregulation and pathway enrichment in *Arid2-deficient* B cells.

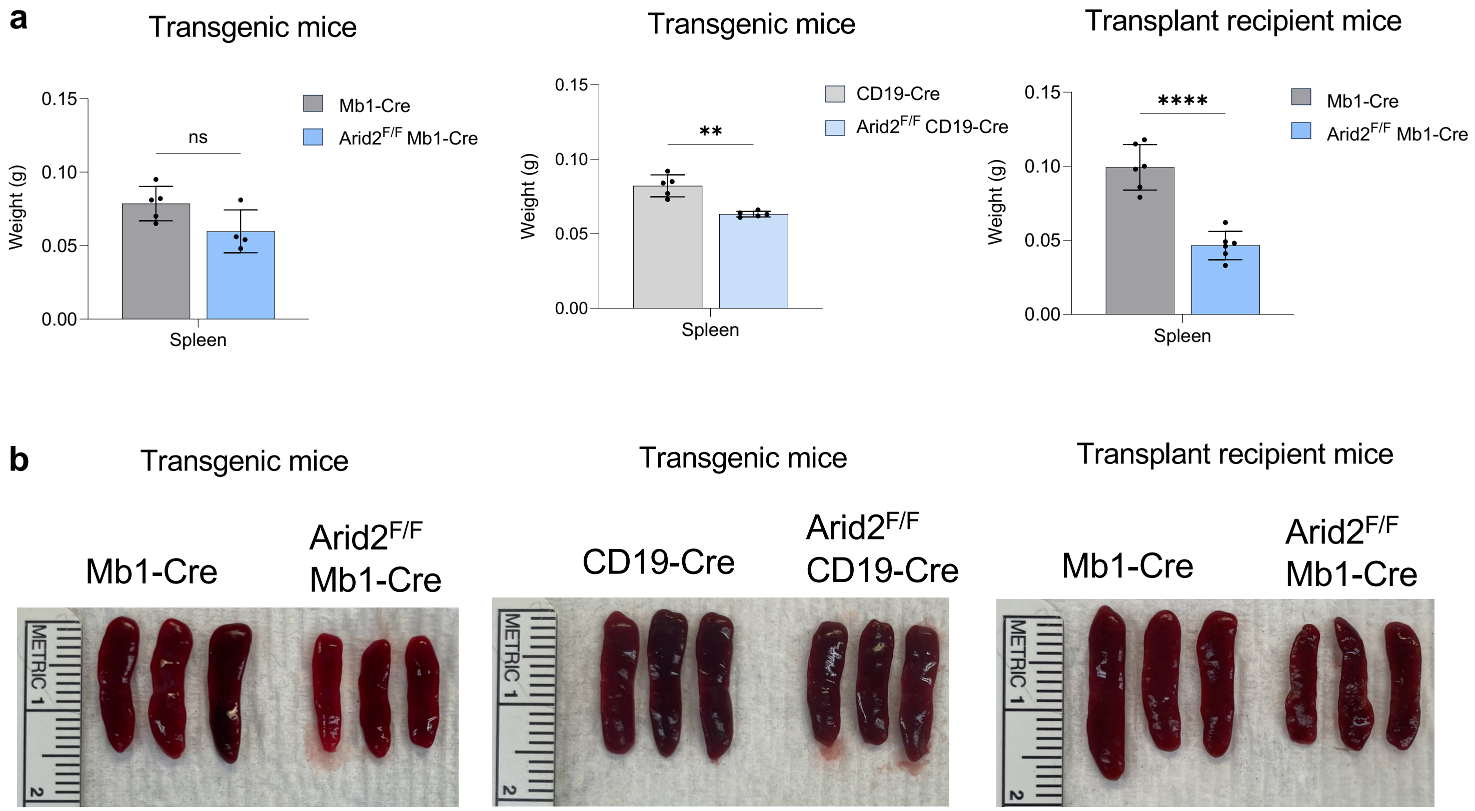


**Figure S4.** Spleen weight and appearance in *Arid2* conditional knockout mice

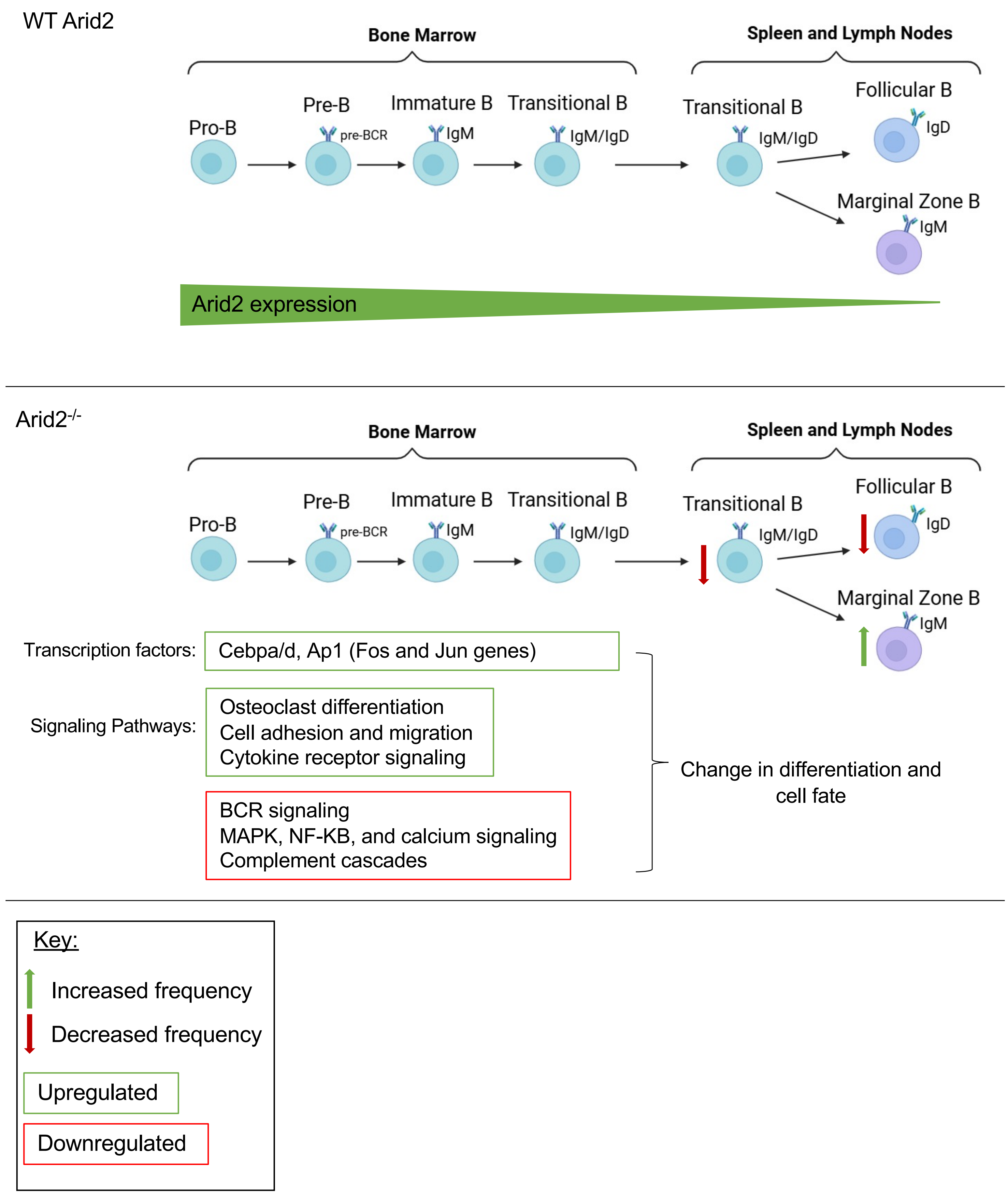
WT Arid2
Bone Marrow
Spleen and Lymph Nodes
Pro-B
Pre-B
pre-BCR
Immature B
IgM
Transitional B
IgM/IgD
Transitional B
IgM/IgD
Follicular B
IgD
Marginal Zone B
IgM
Arid2 expression
Arid2-/-
Bone Marrow
Spleen and Lymph Nodes
Pro-B
Pre-B
pre-BCR
Immature B
IgM
Transitional B
IgM/IgD
Transitional B
IgM/IgD
Follicular B
IgD
Marginal Zone B
IgM
Transcription factors:
Cebpa/d, Ap1 (Fos and Jun genes)
Signaling Pathways:
Osteoclast differentiation
Cell adhesion and migration
Cytokine receptor signaling
BCR signaling
MAPK, NF-KB, and calcium signaling
Complement cascades
Change in differentiation and cell fate
Key:
Increased frequency
Decreased frequency
Upregulated
Downregulated
