## Supplementary material for "Arid2 promotes Follicular B-cell differentiation and antibody responses *in vivo*": Tables

**Table 1: Antibodies used for Flow Cytometry and FACS**

| Antibody Target | Fluorophore | Company | Catalog# | Dilution Factor |
|---|---|---|---|---|
| B220 | PB | BioLegend | 103227 | 1:100 |
| CD23 | BV421 (BrilliantViolet 421) | BioLegend | 101621 | 1:100 |
| CD43 | FITC | BioLegend | 143203 | 1:100 |
| CD21/35 | FITC | BioLegend | 123407 | 1:100 |
| CD3 | FITC | BioLegend | 100203 | 1:100 |
| GL7 | FITC | BioLegend | 144603 | 1:100 |
| CD45.2 | APC | BioLegend | 109814 | 1:100 |
| CD19 | APC | BioLegend | 115512 | 1:100 |
| B220 | APC | BioLegend | 103212 | 1:100 |
| Fas/CD95 | APC | BioLegend | 152604 | 1:100 |
| IgD | AF700 (AlexaFluor 700) | BioLegend | 405730 | 1:100 |
| Gr1 | AF700 (AlexaFluor 700) | BioLegend | 108422 | 1:100 |
| CD19 | PE | BioLegend | 115508 | 1:100 |
| CD93 | PE | BioLegend | 136504 | 1:100 |
| CD43 | PE | BioLegend | 143206 | 1:100 |
| CD3 | PE | BioLegend | 100206 | 1:100 |
| CD43 | PercP-Cyanine5.5 | BioLegend | 143219 | 1:100 |
| CD45 | PercP-Cyanine5.5 | BioLegend | 103132 | 1:100 |
| IgM | PE/Cyanine7 | BioLegend | 406514 | 1:100 |
| Mac1 | PE/Cyanine7 | BioLegend | 101216 | 1:100 |
| CD38 | PE/Cyanine7 | BioLegend | 102717 | 1:100 |
| Ki67 | PE/Cyanine7 | Biolegend | 652426 | 5ul/million cells |
| Annexin V | PE/Cyanine7 | BioLegend | 640949 | 5ul/million cells |
| Live/dead | Propidium iodide (PI) | BioLegend | 421301 | 1:500 |
| Live/dead | 7-AAD | BioLegend | 420403 | 1:500 |

**Table 2: Allele genotyping PCR primers**

| | |
|---|---|
| **Arid2 allele** | |
| Arid2 Forward | 5’-CCTGGCCTTATGGAGGCATT-3’ |
| Arid2 Reverse Floxed | 5’-GCTCTCTGTGTAAGGTGACC-3' |
| Arid2 Reverse Deleted | 5’-GAAATGTAGCAGACTCCACC-3' |
| **Mb1-Cre** | |
| Mb1cre Forward | 5’-CATTTTCGAGGGAGCTTCA-3' |
| Mb1cre Reverse | 5’-ACTGAGGCAGGAGGATTGG-3' |
| **CD19-Cre** | |
| Internal Control Forward | 5’-CTAGGCCACAGAATTGAAAGATCT-3’ |
| Internal Control Reverse | 5’-GTAGGTGGAAATTCTAGCATCATCC-3' |
| CD19cre Forward | 5’-CAGGGTGTTATAAGCAATCCC-3' |
| CD19cre Reverse | 5’- CCTGGAAAATGCTTCTGTCCG-3' |

**Table 3. Surface marker antibody staining**

| **Panel / Context** | **Antibodies Used** |
|---|---|
| **Peripheral blood lineage panel (transgenic mice)** | CD45-PerCP-Cy5.5, B220-PB, CD3-PE, Mac1-PE/Cy7, Gr1-AF700 |
| **Peripheral blood lineage panel (with TdTomato)** | CD45-APC, B220-PB, CD3-FITC, Mac1-PE/Cy7, Gr1-AF700, TdTomato (YL1) |
| **Peripheral blood lineage panel (noncompetitive transplants)** | CD45-APC, B220-AF700, CD3-PE, Mac1-PE/Cy7 |
| **Bone marrow progenitor panel** | B220-PB, IgM-PE/Cy7, CD19-APC, CD43-FITC |
| **Transitional B cell panel** | Bone marrow/PB: B220-APC; Spleen: CD19-APC; plus CD93-PE, CD21/35-FITC, CD23-BV421, IgM-PE/Cy7, IgD-AF700 |

| | |
|---|---|
| **Follicular & MZB panel** | CD19-APC, CD43-PE, CD21/35-FITC, CD23-BV421, IgM-PE/Cy7, IgD-AF700 |
| **Follicular & MZB panel (with TdTomato)** | CD19-APC, CD43-PerCP-Cy5.5, CD21/35-FITC, CD23-BV421, IgM-PE/Cy7, IgD-AF700, TdTomato (YL1) |
| **Germinal Center B cell panel** | CD19-PE, GL7-FITC, Fas/CD95-APC, IgD-AF700, CD38-PE/Cy7 |
| **Follicular & MZB panel with Ki67** | CD19-APC, CD43-PE, CD21/35-FITC, CD23-BV421, Ki67-V |

**Table 4. ImmGen population phenotypes used for gene expression analysis that are shown in Figure 1a.**

| **Short name** | **Full Name** | **Phenotype** | **Location** |
|---|---|---|---|
| proB.CLP.BM | Common Lymphoid Progenitor | Lin-AA4+Kit+IL7Ra+B220- | Bone marrow |
| proB.FrA.BM | Fr. A (pre-pro-B) | Lin-AA4+Kit+IL7Ra+B220+ | Bone marrow |
| proB.FrBC.BM | Fr. B/C (pro-B) | AA4+IgM-CD19+CD43+HSA+ | Bone marrow |
| preB.FrC.BM | Fr. Cprime (cycling pre-B) | AA4+IgM-CD19+CD43+HSA++ | Bone marrow |
| preB.FrD.BM | Fr D (pre-B) | AA4+IgM-CD19+CD43-HSA+ | Bone marrow |
| B.FrE.BM | Fr. E (newly-formed B) | AA4+ IgM+ CD19+ HSA+ | Bone marrow |
| proB.CLP.FL | Common Lymphoid Progenitor | Lin-AA4+Kit+IL7Ra+B220- | Fetal Liver |
| proB.FrA.FL | Fr. A (pre-pro-B) | Lin-AA4+Kit+IL7Ra+B220+ | Fetal Liver |
| proB.FrBC.FL | Fr. B/C (pro-B) | AA4+IgM-CD19+CD43+HSA+ | Fetal Liver |
| preB.FrD.FL | Fr D (pre-B) | AA4+IgM-CD19+CD43-HSA+ | Fetal Liver |
| B.FrE.FL | Fr. E (newly-formed B) | AA4+ IgM+ CD19+ HSA+ | Fetal Liver |
| B.T1.Sp | T1 (transitional) | CD19+B220+IgM++AA4+CD23- | Spleen |
| B.T2.Sp | T2 (transitional) | CD19+B220+IgM++AA4+CD23+ | Spleen |
| B.T3.Sp | T3 (transitional) | CD19+B220+IgM+AA4+CD23+ | Spleen |
| B.Fo.Sp | Fo (follicuilar) | CD19+ B220+ IgM+ AA4- CD23+ | Spleen |
| B.GC.Sp | Germinal Center B cells, Spleen | CD19+IgM+IgD-GL7+PNA+ | Spleen |
| B.MZ.Sp | MZ (Marginal Zone) | CD19+ B220+ IgM+ AA4- CD23- CD21/35+ | Spleen |
| B1a.Sp | B-1a | CD19+ B220+ IgM+ AA4- CD23- CD43+ | Spleen |
| B.FrF.BM | Fr. F (recirc B) | AA4- IgM+ CD19+ | Bone marrow |
| B.Fo.MLN | Mesenteric node B cells | CD19+ B220+ IgMdull AA4- CD23+ CD43- CD21/35+ | Mesenteric Lymph Node |

| | | | |
|---|---|---|---|
| B.Fo.LN | Lymph node B cells | CD19+ B220+ IgMdull AA4- CD23+ CD43- | Lymph Node |
| B.Fo.PC | Fo-type, peritoneal cavity | CD19+ IgM+ CD23+ CD43- CD5- | Peritoneal Cavity |
| B1b.PC | B-1 b, peritoneal cavity | CD19+ IgM++ CD43+ CD5- | Peritoneal Cavity |
| B1a.PC | B-1a, peritoneal cavity | CD19+ IgM++ CD43+ CD5+ | Peritoneal Cavity |
